# Single-cell and spatial detection of senescent cells using DeepScence

**DOI:** 10.1101/2023.11.21.568150

**Authors:** Yilong Qu, Beijie Ji, Runze Dong, Liangcai Gu, Cliburn Chan, Jichun Xie, Carolyn Glass, Xiao-Fan Wang, Andrew B Nixon, Zhicheng Ji

## Abstract

Accurately identifying senescent cells is essential for studying their spatial and molecular features. We developed DeepScence, a method based on deep neural networks, to identify senescent cells in single-cell and spatial transcriptomics data. DeepScence is based on CoreScence, a senescence-associated gene set we curated that incorporates information from multiple published gene sets. We demonstrate that DeepScence can accurately identify senescent cells in single-cell gene expression data collected both *in vitro* and *in vivo*, as well as in spatial transcriptomics data generated by different platforms, substantially outperforming existing methods.

## Introductions

Cellular senescence is a fundamental biological process in which cells enter a state of permanent cell cycle arrest, losing their ability to proliferate [1, 2]. This state primarily serves as a defense mechanism against the uncontrolled cell division observed in cancer. However, the accumulation of senescent cells (SnCs) over time may contribute to the decline in the physical and functional integrity of tissues and organs, playing a significant role in aging [3, 4]. Additionally, SnCs are implicated in the development of various age-related diseases, such as osteoarthritis, pulmonary fibrosis, and Alzheimer’s disease [5–8]. SnCs secrete pro-inflammatory proteins that alter the cellular microenvironment, affecting neighboring cells and contributing to chronic inflammation [9]. Given its profound impact on human health and its potential as a therapeutic target, it is important to understand cellular senescence and identify its associated biomarkers.

Given the relatively rare presence of SnCs in tissues [10], technologies capable of achieving single-cell or near single-cell resolution, such as single-cell RNA sequencing (scRNA-seq) and spatial transcriptomics (ST), have shown great promise in uncovering the molecular and spatial features of SnCs [10–12]. Several methods have been developed to identify SnCs in scRNA-seq data. However, their reliability remains questionable due to various challenges. The first type of method uses a single gene marker, such as CDKN1A (the gene that encodes the p21 protein) or CDKN2A (the gene that encodes the p16 protein), to identify SnCs[13, 14]. If the gene shows positive expression in a cell, the cell is identified as an SnC. However, an SnC may not exhibit positive expression of the marker gene due to dropout events [15], and the results of such methods can be highly unreliable due to the high levels of noise and sparsity in scRNA-seq data.

The second type of method scores the senescence level of a cell by ranking the genes in a senescence gene set (SnG) using methods such as AUCell and ssGSEA [16, 17]. However, these methods lack the ability to capture the complex nonlinear and combinatorial relationships between a cell’s senescence level and the expression of genes in the SnG. Furthermore, identifying a reliable SnG is challenging, and we will demonstrate in this study that there is very little agreement across published SnGs. Additionally, genes in an SnG can either induce or inhibit senescence, but this directionality is ignored by these methods.

A recently published method, SenCID [18], is the third type of method that trains supervised machine learning models to predict SnCs. SenCID trains a support vector machine (SVM) using published bulk RNA-seq data from both normal and senescent cells. The trained model can predict SnCs in both single-cell and bulk RNA-seq datasets. However, a major limitation of SenCID is that its training data were collected from *in vitro* studies. It is known that the molecular characteristics of cellular senescence can differ between *in vitro* and *in vivo* conditions[19]. Therefore, it is questionable whether the trained SenCID models can reliably predict SnCs in *in vivo* datasets. Additionally, trained SenCID models may not be directly applicable to datasets generated by ST technologies, such as 10x Xenium, which have different data distributions and number of profiled genes compared to scRNA-seq.

To address these issues, we developed DeepScence, an unsupervised machine learning model based on an autoencoder architecture for identifying SnCs. We designed a customized autoencoder that leverages CoreScence, a core senescence gene set we compiled, to efficiently capture senescence-related information. We systematically evaluated the performance of DeepScence and found that it accurately identifies SnCs in both *in vivo* and *in vitro* settings, as well as in both scRNA-seq and ST datasets. DeepScence substantially outperforms the three types of existing methods, including SenCID. DeepScence paves the way for further studies on the spatial and molecular features of SnCs in various contexts.

## Results

### Large discrepancies across existing SnGs

A reliable senescence gene set (SnG) is crucial for identifying senescent cells (SnCs). We surveyed nine published SnGs, including SenMayo [20], SenSig [21], CSGene [22], SeneQuest [2], CellAge [23], GenAge [24], SASP-related genes from De Cecco et al. [25], inflammatory network genes in senescence from Freund et al. [9], and the transcriptome signature of senescence from Casella et al. [26]. Three gene sets (SenSig, De Cecco et al., and Casella et al.) were obtained from data-driven approaches, such as differential analysis between normal and senescent cells using RNA-seq data. The other six gene sets were obtained through literature searches. Strikingly, we found a large discrepancy across these SnGs. First, the number of genes included in each SnG varies greatly (Figure 1a). Larger SnGs, such as SenSig, contain more than a thousand genes, whereas smaller ones, like the Casella et al. SnG, include fewer than a hundred genes. Second, there is an extremely low level of agreement across SnGs (Figure 1b). The Jaccard index, which measures the degree of overlap between two sets, is below 0.2 for nearly all SnG pairs. Third, a gene reported by one SnG may not be consistently found in another. Out of the 2,966 senescence-related genes identified by at least one SnG, 2,052 (69.2%) were reported in only one SnG, and only 39 (1.3%) were reported in at least five SnGs (Figure 1c).

**Figure 1.**
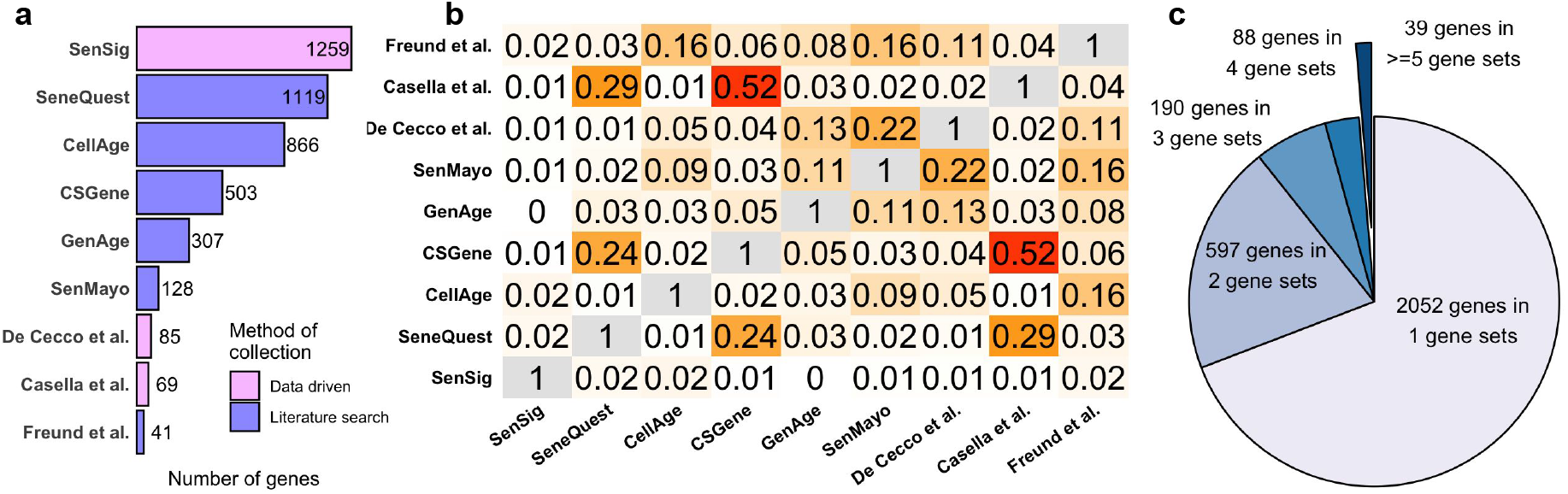
**a**, Number of genes in each SnG. Colors represent the two different methods by which each gene set was derived. **b**, Jaccard index between each pair of SnGs. **c**, Number of genes reported by different numbers of SnGs.

Since these SnGs were compiled by different research groups, the discrepancies could stem from variations in scope and inclusion criteria. As a result, SnC identification methods that rely on a single SnG may produce biased and irreproducible outcomes.

### CoreScence: a core senescence gene set

To address this discrepancy, we compiled a new senescence gene set, CoreScence, which consists of 39 genes reported by at least five published gene sets (Figure 2a, Supplementary Table S1). The rationale is that genes consistently reported by multiple gene sets are less likely to be influenced by inclusion bias and are more likely to be associated with senescence. We found that using a cutoff of five published gene sets often yields better performance on real data (Supplementary Figure 1). CoreScence includes several canonical marker genes for cellular senescence, such as CDKN1A and CDKN2A [1, 10]. It is important to note that, in addition to senescence, genes in CoreScence may also be involved in other functions or pathways, such as immunity or development. This will be accounted for in the DeepScence model described below.

**Figure 2.**
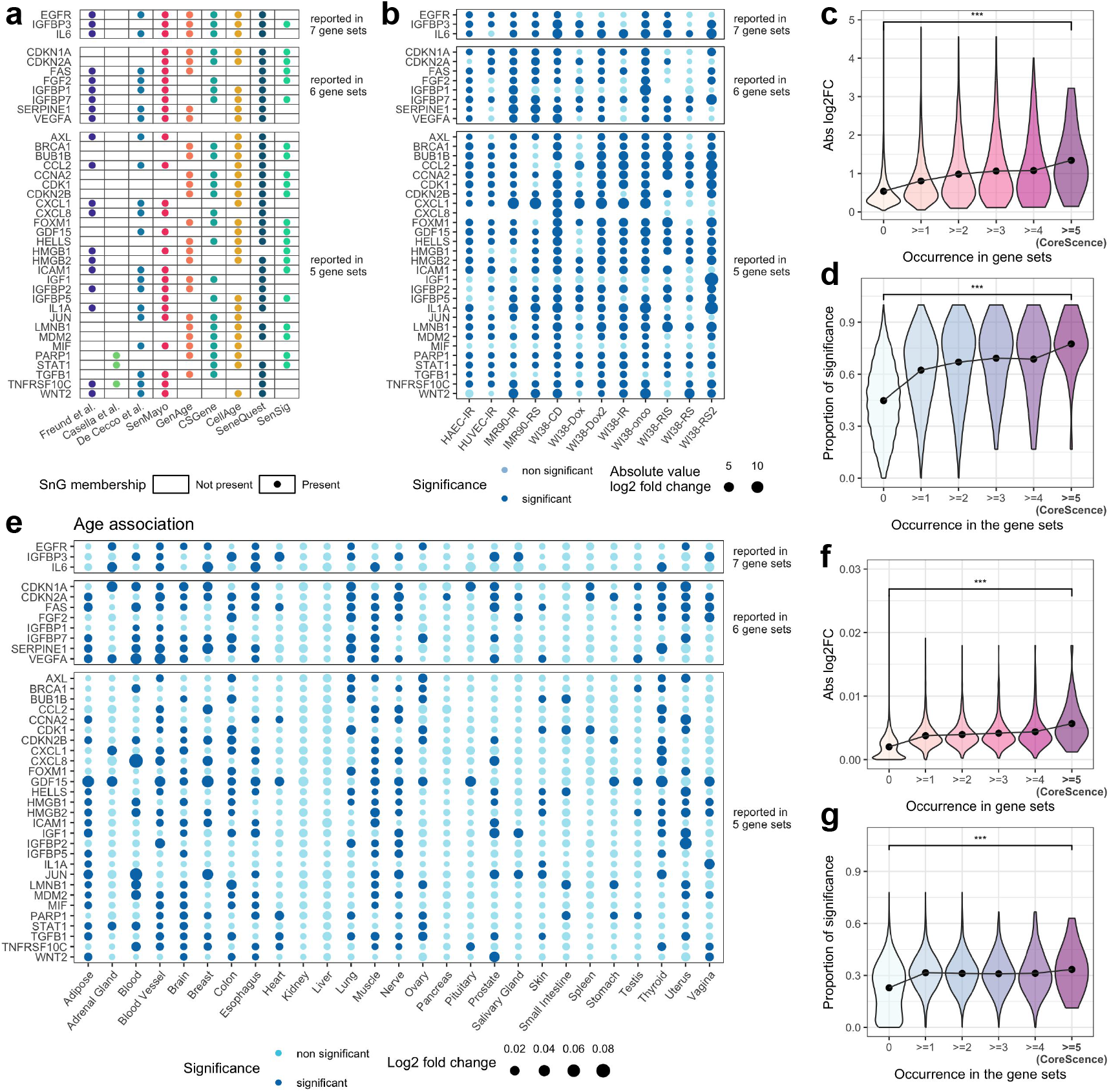
**a**, The CoreScence genes and their SnG memberships. **b**, Differential expression in bulk RNA-seq data for the CoreScence genes. Colors indicate statistical significance, and the sizes of the dots represent the absolute values of log_2_ fold change. Column names indicate the cell line and senescence induction methods. IR refers to ionizing radiation; RS refers to replicative senescence; CD refers to cell density; Dox refers to Doxorubicin; RIS refers to Ras-induced senescence; onco refers to oncogene-induced senescence. **c**, Averaged absolute values of log_2_ fold change across DEG comparisons (y-axis) and the number of SnGs reporting a gene (x-axis). Wilcoxon test was conducted to compare the two distributions. “***” indicates p-value < 0.001. Black dots represent the mean values. **d**, Proportion of DEG comparisons with statistically significant differential expression (y-axis) and the number of SnGs reporting a gene (x-axis). Wilcoxon test was conducted to compare the two distributions. “***” indicates p-value < 0.001. Black dots represent the mean values. **e**. Differential expression of CoreScence genes in GTEx bulk RNA-seq data across age. Colors indicate statistical significance, and dot sizes represent the absolute values of the log_2_ fold change. Column names indicate the tissues from which the data were obtained. **f**. Average absolute log_2_ fold change values across age-related DEG comparisons (y-axis) plotted against the number of SnGs reporting a gene (x-axis). A Wilcoxon test was conducted to compare the two distributions. “***” indicates *p <* 0.001. Black dots represent mean values. **g**. Proportion of DEG comparisons with statistically significant differential expression (y-axis) plotted against the number of SnGs reporting a gene (x-axis). A Wilcoxon test was conducted to compare the two distributions. “***” indicates *p <* 0.001.

To validate CoreScence, we collected information on genes with differential expression between senescent and non-senescent cells across various cell lines from published bulk RNA-seq studies [26, 27] (Figure 2b). We found that genes consistently reported by a greater number of published SnGs exhibit stronger differential signals between senescent and non-senescent cells (Figure 2c-d). In addition, we found that genes reported by a greater number of published SnGs are more likely to be associated with sample age (Figure 2e–g). These results suggest that the genes in CoreScence are more likely to be genuinely associated with senescence.

### DeepScence: a deep neural network for identifying SnCs

Building upon CoreScence, we developed DeepScence, a machine learning model based on an autoencoder architecture for identifying SnCs (Figure 3a, Methods). Autoencoders have been successfully applied to scRNA-seq analysis in previous studies[28–30], demonstrating their ability to capture complex, combinatorial, and non-linear patterns in the data. As an unsupervised method, autoencoders also offer greater flexibility and are better suited to generalize across diverse biological contexts. The input to DeepScence is a gene expression count matrix of the genes in the CoreScence gene set, and we assume that these counts follow zero-inflated negative binomial (ZINB) distributions [31]. The bottleneck layer of the autoencoder is designed to consist of two neurons that are almost uncorrelated. One neuron is responsible for capturing senescence-related information, while the other captures information unrelated to senescence. The output of DeepScence consists of three components corresponding to the three parameters of the ZINB distribution. After model fitting, the continuous value of the neuron capturing senescence information, referred to as the senescence score, is the final output of DeepScence. The continuous senescence scores can be optionally binarized through a permutation-based procedure to classify cells as either senescent or non-senescent (Methods).

**Figure 3.**
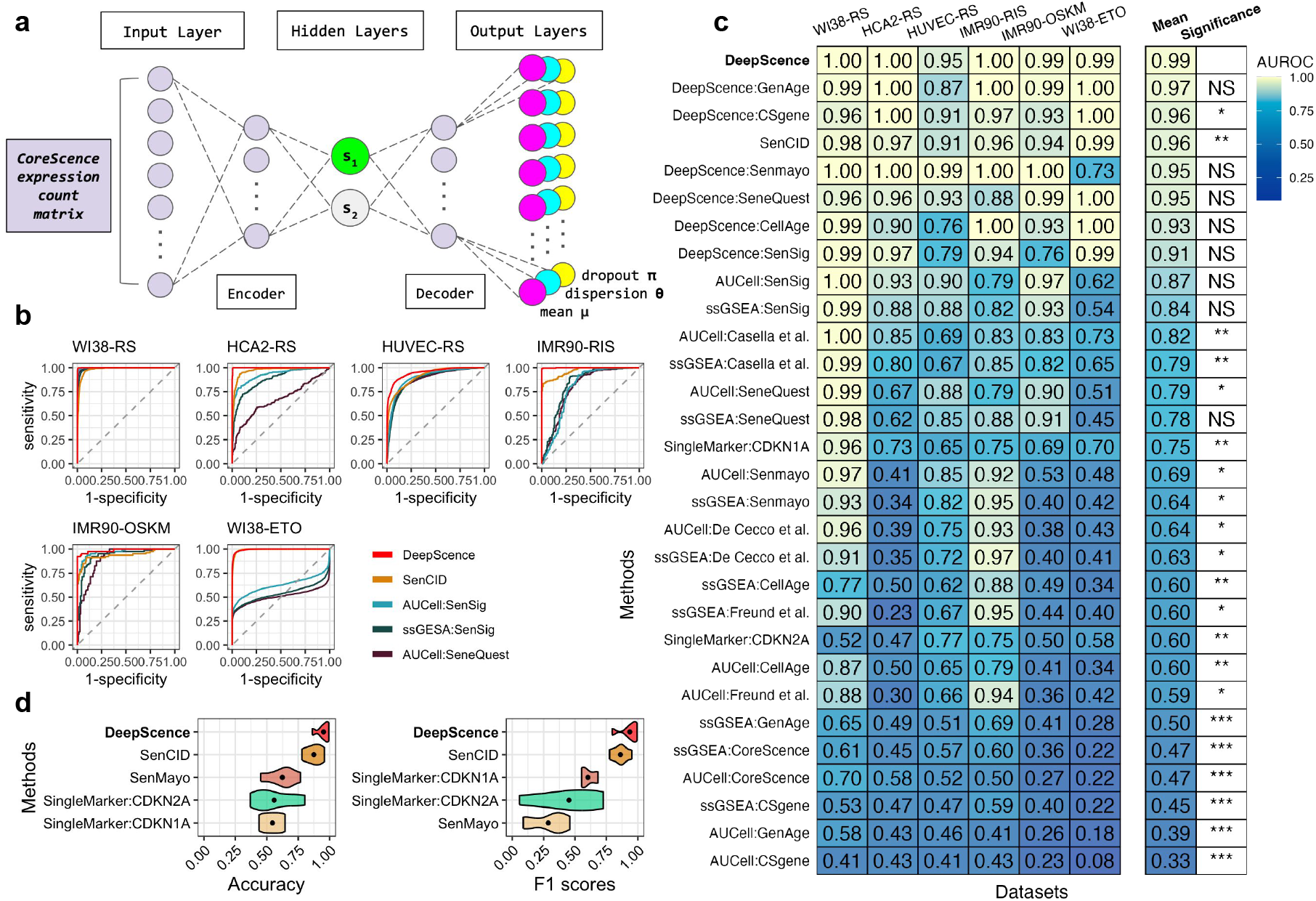
**a**, DeepScence model architecture. The green neuron, s1, in the bottleneck layer represents the neuron that captures senescence score. **b**, ROC curves of top-performing methods across *in vitro* datasets. **c**, AUROCs for all methods across *in vitro* datasets. Methods are ordered in decreasing order by average AUROCs. Paired t-test was conducted to compare the performance between DeepScence and each of the other methods. “***” indicates p-value < 0.001, “**” indicates 0.001 < p-value < 0.01, “*” indicates 0.01 < p-value < 0.05, and “NS” indicates not significant (p-value > 0.05). **d**, Accuracy and F1 scores across *in vitro* datasets, comparing DeepScence with existing methods capable of binarization. Dots in the violin plots represent mean values.

### DeepScence accurately identifies SnCs in *in vitro* scRNA-seq datasets

We applied DeepScence and three existing types of SnC identification methods (Methods) to six *in vitro* studies [27, 32–36]. We also evaluated variants of DeepScence in which published gene sets were used instead of CoreScence as input to the same autoencoder model. In each study, scRNA-seq was performed on cell lines both before and after senescence induction (Supplementary Table S2). We first evaluated the performance of the continuous senescence scores generated by each method, using senescence induction as the gold standard. Figure 3b shows the receiver operating characteristic (ROC) curves for the top-performing methods in each dataset, and Figure 3c shows the area under the ROC curve (AUROC) as an evaluation metric. Both DeepScence and SenCID are among the best performing methods, with AUROCs exceeding 0.9 across all datasets. DeepScence exhibited slightly better performance than SenCID in every dataset, and showed significantly better overall performance. The variants of DeepScence also show strong performance, indicating that DeepScence is robust to variations in the input gene sets and that the autoencoder model is the main contributor to the improved performance. Nevertheless, the default DeepScence with CoreScence as input still achieves the best performance, suggesting that CoreScence comprises genes that are particularly relevant for predicting SnCs. For some SnGs, gene ranking-based methods performed well overall but failed in certain datasets with AUROCs below 0.8, potentially due to their limited ability to capture combinatorial and nonlinear relationships in gene expression. Other SnGs showed poor performance using these methods. Single marker gene-based SnC identification methods also performed poorly, potentially due to the high sparsity of scRNA-seq data and the limited information conveyed by a single gene.

For each method, we further classified cells into two discrete categories, senescent and non-senescent, by binarizing the continuous senescence scores (Methods). We evaluated the performance of these binarized groups using accuracy and F1 scores (Figure 3d). Once again, DeepScence and SenCID were the top-performing methods, with DeepScence slightly outperforming SenCID. The other methods showed substantially worse performance in both accuracy and F1 scores.

### DeepScence outperforms existing methods in *in vivo* scRNA-seq datasets

We next evaluated all methods for identifying SnCs in *in vivo* scRNA-seq datasets. We collected eight datasets in which scRNA-seq was performed on human and mouse tissues *in vivo* [37–43]. These datasets cover various tissue types and disease models, such as autoimmune orchitis, muscle injury, lung idiopathic pulmonary fibrosis (IPF), gingival senescence, heart pressure overload, Alzheimer’s disease (AD) in the brain, and lung cancer. Confirmed by immunohistochemistry staining (Supplementary Table S2), these studies found that senescent cells were more enriched in certain contexts, such as fibrosis, for specific cell types. We again used AUROC as a metric to assess the performance of each method.

Figure 4a shows the overall performance of each method. Figure 4b shows the ROC plots for methods that performed best in the *in vitro* datasets. DeepScence and its variants are the only methods that have reliable overall performance, with an average AUROC of 0.86 for DeepScence. The averaged AUROC of all other methods drop significantly, with the second best method reaching only 0.77. Moreover, methods other than DeepScence fail in at least one cell type with AUROC smaller than 0.6. SenCID, that has a performance comparable to DeepScence in *in vitro* datasets, performs poorly in *in vivo* datasets, with AUROC less than or equal to 0.2 in five cell types. The reason is that SenCID assigns substantially lower senescence scores to the cell population with enriched senescent cells, contradictory with experimental findings. In comparison, the distributions of senescence scores assigned by DeepScence agree with the enrichment of senescent cells in all cases (Figure 4c).

**Figure 4.**
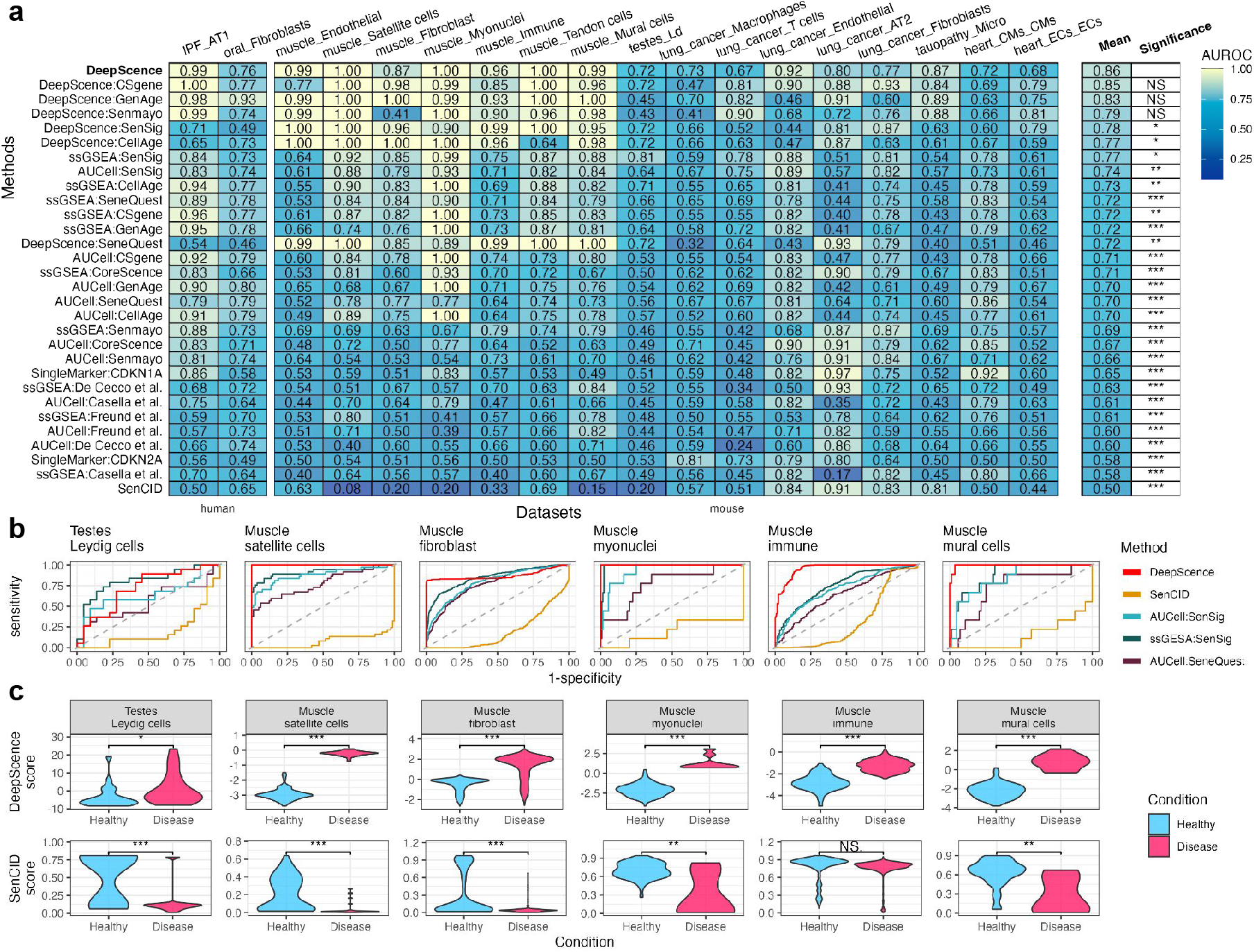
**a**, AUROCs for all methods across *in vivo* datasets. Methods are ordered in decreasing order by average AUROCs. Paired t-test was conducted to compare the performance between DeepScence and each of the other methods. “***” indicates p-value < 0.001, “**” indicates 0.001 < p-value < 0.01, “*” indicates 0.01 < p-value < 0.05, and “NS” indicates not significant (p-value > 0.05). **b**, ROC curves for six example cell types in *in vivo* datasets, showing methods with top performance in *in vitro* datasets. **c**, Distribution of SenCID and DeepScence scores, comparing cells from healthy and diseased conditions. Wilcoxon test was conducted to compare the two distributions in each case. ‘*’ indicates p-value between 0.01 and 0.05, ‘**’ indicates p-value between 0.001 and 0.01, and ‘***’ indicates p-value < 0.001.

### DeepScence outperforms existing methods in spatial transcriptomics datasets

We further applied DeepScence and other competing methods to two ST datasets. The first dataset includes ST samples of mouse muscle generated by 10x Visium [38, 44]. The tissue slices were subjected to injury induced by notexin, and ST data were collected at 2 and 5 days post-injury. The injured regions were distinguishable from healthy regions by histology (Figure 5a), showing higher *β*-gal intensities and distinct cell morphology (Figure 5b), indicating that SnCs were more enriched in the injured regions. The second dataset comprises ST samples from human brains diagnosed with severe Alzheimer’s disease (AD), along with age-matched, normally aging controls. This dataset was generated using Stereo-seq, an ST platform with subcellular spatial resolution, enabling the identification and extraction of single cells for downstream analysis. Prior studies [41, 45] have reported that microglial senescence is more prevalent in AD than in non-diseased brains, supported by increased immunostaining of senescence markers such as *β*-galactosidase, p16, p21 [46], and *γ*H2AX [47].

**Figure 5.**
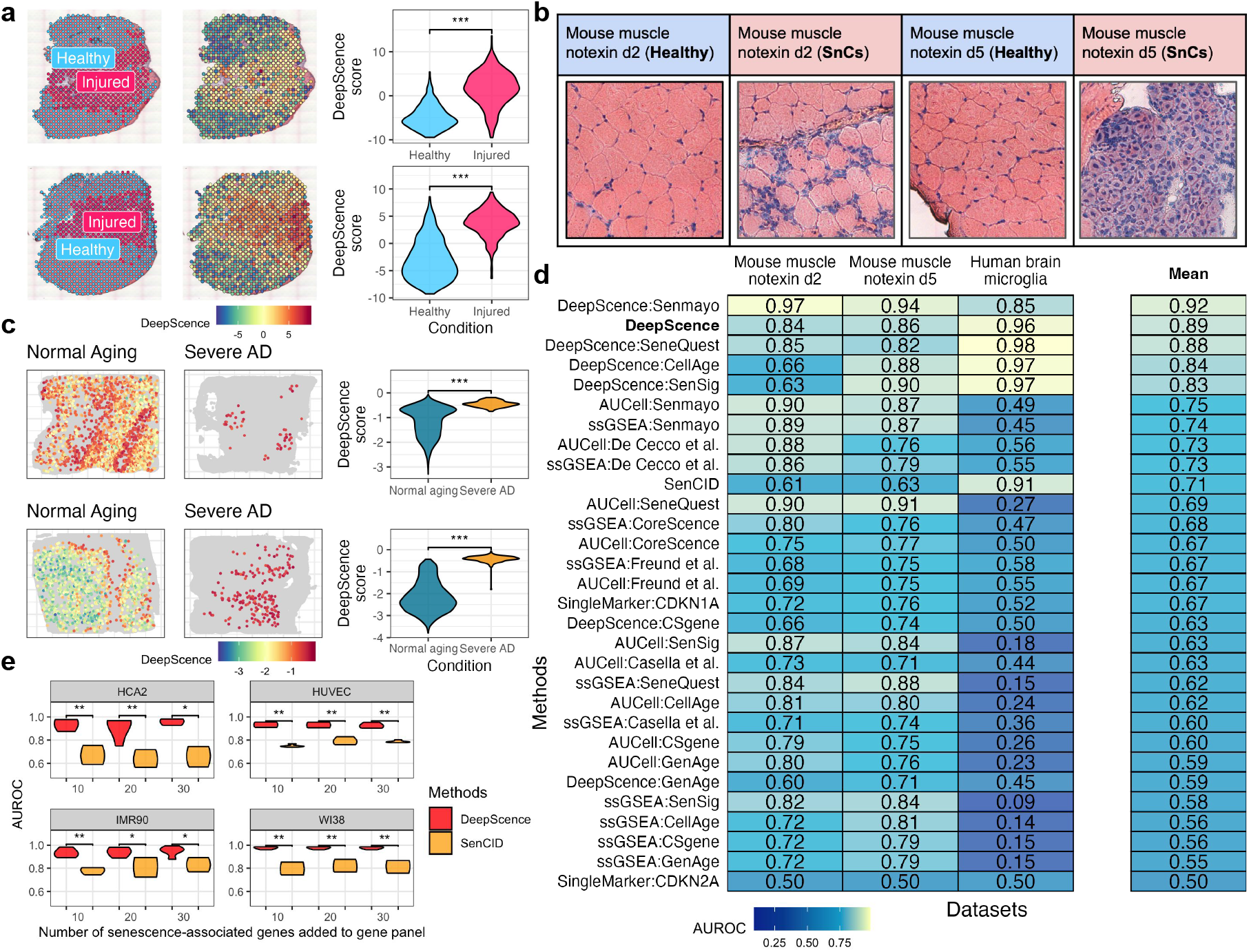
**a**, From left to right: spatial spots colored by healthy and injured regions, spatial spots colored by DeepScence scores, and the distribution of DeepScence scores comparing the two conditions. The Wilcoxon test was used to compare the two distributions within each sample. “***” indicates a p-value < 0.001. The top and bottom rows represent muscle samples at 2 and 5 days post-injury, respectively. **b**, Enlarged H&E images showing cell morphology in SnC and healthy regions for each of the two muscle samples. **c**, From left to right: spatial locations of microglia colored by DeepScence scores, with other cells shown in grey, in normal aging and severe AD samples, and the distribution of DeepScence scores comparing the two conditions. The Wilcoxon test was used to compare the distributions in each sample. “***” indicates a p-value < 0.001. **d**, AUROCs of all methods across ST datasets. Methods are ordered in decreasing order by their average AUROC. **e**, AUROC (y-axis) of DeepScence and SenCID in simulated Xenium datasets with different numbers of senescence-associated genes added to the default 10x gene panel (x-axis). Wilcoxon test was conducted to compare DeepScence and SenCID AUROCs. ‘*’ indicates p-value between 0.01 and 0.05, ‘**’ indicates p-value between 0.001 and 0.01, and ‘***’ indicates p-value < 0.001.

DeepScence assigned higher senescence scores to Visium spots in the injured regions compared to the healthy regions in the first dataset (Figure 5a), and to microglial cells from severe AD patients compared to those from normally aging individuals (Figure 5c). DeepScence and its variants are among the best-performing methods across both datasets, substantially outperforming other competing methods (Figure 5d). These examples further demonstrate DeepScence’s superior performance in both *in vivo* and spatial settings.

We also applied DeepScence and SenCID, the two top-performing methods in *in vitro* datasets, to synthetic 10x Xenium datasets. Unlike 10x Visium, which profiles nearly the entire transcriptome, 10x Xenium, similar to other ST technologies based on fluorescence in situ hybridization (FISH), can only profile hundreds of prespecified genes [48]. For evaluation purposes, we synthesized 10x Xenium datasets from *in vitro* scRNA-seq data by subsetting a number of genes comparable to actual 10x Xenium data, while adding a small number of senescence-associated genes to the default 10x gene panel (Methods).

DeepScence performs consistently well across different datasets and with varying numbers of genes in the gene panel, whereas SenCID’s performance drops considerably (Figure 5e). These results demonstrate the generalizability of DeepScence to data types beyond scRNA-seq. Moreover, DeepScence requires the addition of only 10 senescence-associated genes to achieve strong performance, allowing for the exploration of other biological pathways and functions during gene panel design.

### Analyzing normally aging tissues with DeepScence

Finally, we applied DeepScence and other competing methods to scRNA-seq data from the Tabula Sapiens (TS) study [49]. The TS dataset includes various tissues and cell types collected from normal human individuals. Unlike the previously analyzed datasets, where external validation information such as *β*-gal staining was available to establish a gold standard, the TS dataset does not provide direct gold standards for evaluating SnC identification methods. Instead, we assessed whether cells with the highest senescence scores were more likely to originate from older individuals (Methods), based on previous findings that senescent cells accumulate with age and are rare in normal individuals[2, 4, 10].

In 77.61% of the 67 tissue–cell type pairs, DeepScence identifies a higher enrichment of cells from older individuals compared to younger individuals among cells with the highest senescence scores (Figure 6a). DeepScence and its variant are among the methods that yield the highest average enrichment scores across all tissue–cell type pairs (Figure 6b). These results suggest that DeepScence produces senescence scores that better align with existing findings linking senescence to the aging process.

**Figure 6.**
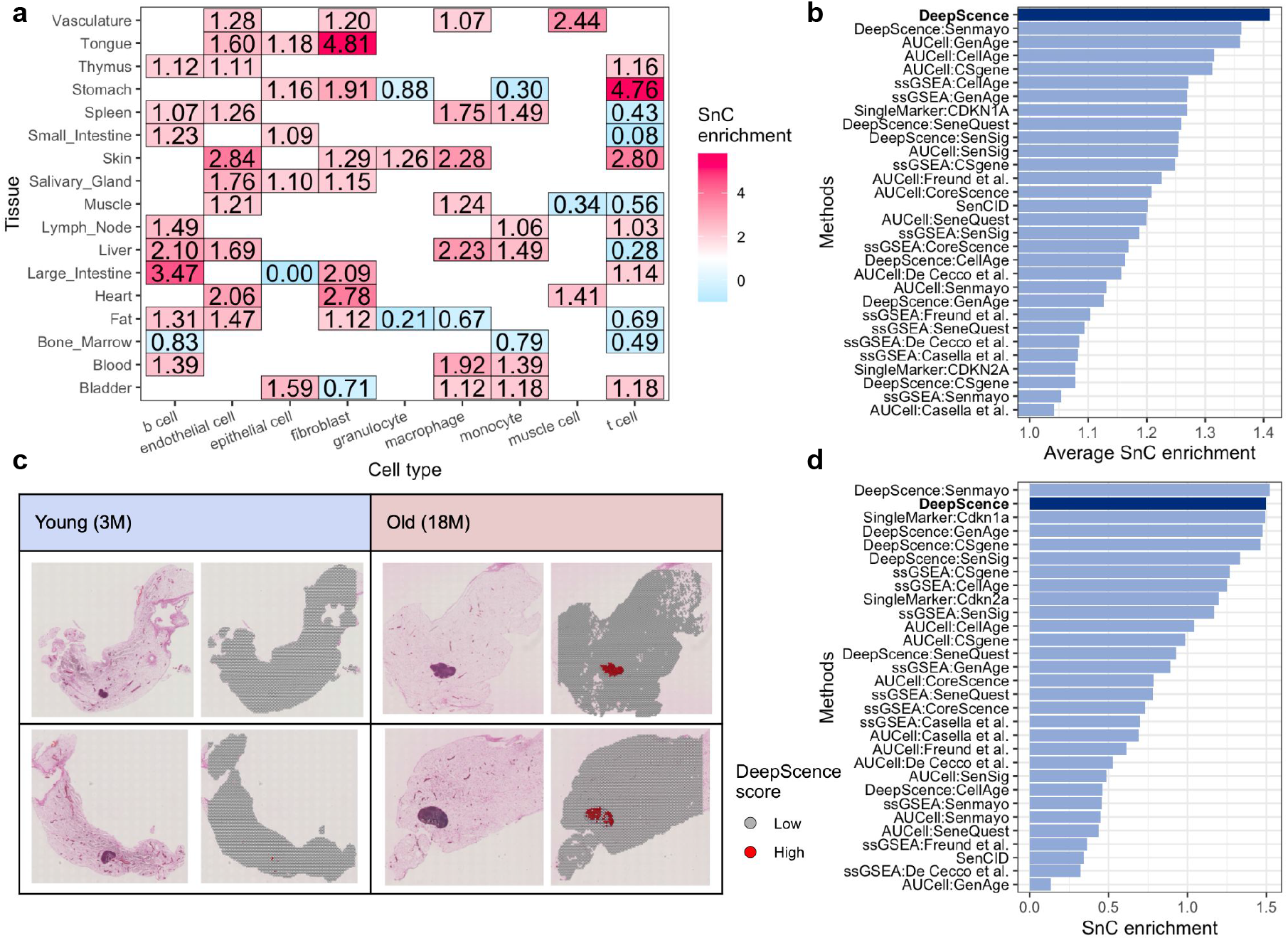
**a**, SnC enrichment scores from DeepScence across tissue–cell type pairs in the Tabula Sapiens (TS) dataset. Red indicates that SnCs identified by DeepScence are more likely from older samples, while blue indicates a higher likelihood of originating from younger samples. **b**, Average SnC enrichment scores for all methods across all tissue–cell type pairs in the TS dataset. **c**, Each panel shows H&E images (left) and the spatial distribution of DeepScence scores (right) for Visium ST samples from young and old mouse mammary glands. Each row represents either a young or an old sample. **d**, SnC enrichment scores for all methods in the mouse mammary gland ST dataset.

We also applied DeepScence and other competing methods to a Visium ST dataset with normally aging mouse mammary gland tissues. The dataset includes two samples from 3-month-old mice and two samples from 18-month-old mice. Almost all spatial spots with the highest DeepScence scores are in the older mice (Figure 6c). Spatial spots with high senescence scores are all enriched in the lymph node region of the two old samples, showing a highly consistent spatial pattern. This is consistent with previous findings [50, 51], which report increased immune cell presence and enhanced epithelial–immune interactions in aged mice. We further numerically evaluated SnC enrichment across old and normal samples similar to the TS dataset (Figure 6d). DeepScence and its variant are still among the best performing methods.

## Conclusions

In this study, we developed DeepScence, a deep-learning framework for identifying senescent cells in single-cell and spatial transcriptomics data. DeepScence is built upon CoreScence, a new senescence gene set we compiled to resolve the large discrepancies across existing gene sets. To evaluate the performance of DeepScence and other competing methods, we collected benchmarking scRNA-seq and ST datasets representing a wide variety of tissue types and disease statuses. In most of the benchmarking datasets, cellular senescence was validated using external information independent of gene expression, such as immunohistochemistry staining. We demonstrated that DeepScence accurately identifies senescent cells in both *in vitro* and *in vivo* scRNA-seq datasets, as well as in ST data generated by 10x Visium and 10x Xenium. Built upon CoreScence, which includes genes generally associated with senescence, DeepScence is able to generalize across various species, tissue types, and disease states. In contrast, existing methods perform poorly in some scenarios. DeepScence is highly flexible and, in principle, can be applied to other types of data, such as proteomics. For research projects that involve different types of data, using DeepScence to detect SnCs not only streamlines the data analysis process but also ensures consistency in analytical approaches throughout the study. DeepScence is also computationally efficient and can scale to large datasets (Supplementary Figure 2). For example, DeepScence can analyze a dataset with 10^5^ cells in 10 minutes, using a maximum of 35.02 GB of memory on an NVIDIA RTX 5000 GPU.

Although DeepScence is built upon a deep neural network, the number of learnable parameters is small due to the limited number of neurons in each hidden layer and the relatively simple model architecture, which relies solely on fully connected layers. For example, for 39 input genes, DeepScence involves only 5303 learnable parameters. This number is substantially smaller than the millions or even billions of parameters required by other deep learning architectures based on convolutional layers or transformers, such as AlexNet[52], ResNet[53], and GPT-3[54]. As a result, DeepScence is less prone to overfitting, even when the number of input genes or cells is relatively small. To further prevent overfitting, we applied a cross-validation procedure in which 10% of the cells were used as a validation set, and the model’s loss was monitored on this set. Additionally, we incorporated an early stopping mechanism to halt training if the validation loss did not improve. These measures enhance the robustness and generalizability of the DeepScence model.

Existing methods for experimentally identifying SnCs have largely relied on techniques such as *β*-gal staining. While vast amounts of single-cell and spatial sequencing data have been generated across various tissue types and disease states, the majority of these datasets lack accompanying *β*-gal staining or other relevant annotations, making gene expression the only available information for identifying SnCs. DeepScence’s ability to accurately identify SnCs based solely on gene expression would fully unlock the potential of these datasets for senescence research. Mapping SnCs in previously unstudied tissues and disease contexts using DeepScence could substantially enhance our understanding of cellular senescence and its roles in diverse diseases, including cancer and age-related conditions, thereby opening new avenues for clinical diagnostics and therapeutic strategies.

## Limitations

The CoreScence gene set, upon which DeepScence is built, consists of genes that are generally associated with senescence and are invariant across specific tissues or cell types. Since signatures of cellular senescence can vary across tissues and cell types, collecting SnGs that are tissue- and cell-type-specific has the potential to further improve the performance of DeepScence. While DeepScence has been developed and tested only on datasets with gene expression information, in principle, it can be extended to handle data from other modalities, such as protein abundances and chromatin accessibility, which is worth exploring in the future. Finally, healthy and senescent cells can often be distinguished by their morphological features. Although DeepScence currently utilizes only molecular information, combining imaging data with molecular information may further enhance its performance.

## Methods

### Construction and evaluation of CoreScence

#### Collection of published SnGs

We collected nine senescence gene sets from published studies. SenMayo was collected from Saul et al. [20], CSGene from Zhao et al. [22], SenSig from Cherry et al. [21], and SASP genes from De Cecco et al. [25]. The gene set for the inflammatory network in cellular senescence was collected from Freund et al. [9], and the transcriptome signature of cellular senescence was collected from Casella et al. [26]. CellAge and GenAge were obtained from the Human Ageing Genomic Resources (HAGR) [23, 24]. The SeneQuest gene set was downloaded from the SeneQuest database [2], and a gene was retained either if it was reported by at least 15 publications, or if it was reported by at least four publications with at least 70% agreement on whether the gene induces or inhibits senescence. This criterion ensures that both frequently reported genes and genes with clear directionality are included.

#### Collection of differentially expressed genes from bulk RNA-seq datasets

We collected information on genes with differential expression between senescent and non-senescent cells from two published bulk RNA-seq datasets. The differential gene data, provided by the original studies, were directly downloaded from the Gene Expression Omnibus (GEO) under accessions GSE130727 and GSE175533. The downloaded data include log fold changes, p-values, and p-values adjusted for multiple testing. The first study [26] (GSE130727) includes HEAC, HUVEC, IMR-90, and WI-38 cell lines, with senescence induced by doxorubicin treatment, ionized radiation, oncogene induction, and replicative senescence via the Hayflick limit. The second study [27] (GSE175533) includes the WI-38 cell line, where senescence was induced by Ras, replicative senescence via the Hayflick limit, and increased cell density.

#### Collection of bulk RNA-seq datasets from normally aging tissues

Gene expression matrices from the Genotype-Tissue Expression (GTEx)[55] bulk RNA-seq dataset were downloaded from the GTEx portal. Tissues with fewer than 100 samples were excluded. Age information for each sample, provided as an interval by GTEx, was converted into a numerical value by averaging the two endpoints of the interval. Within each tissue, genes with log_2_(TPM + 1) values greater than 1 in at least 10% of the samples were retained. Linear regression was performed within each tissue using the R package limma (version 3.54.2)[56]. The response variables were the raw gene expression counts, and the independent variables were age and sex. P-values were adjusted for multiple testing using the Benjamini–Hochberg (BH)[57] procedure, and adjusted p-values less than 0.05 were considered statistically significant.

### DeepScence model

DeepScence utilizes a Zero-Inflated Negative Binomial (ZINB) autoencoder to learn meaningful low-dimensional representations of scRNA-seq datasets and to score each cell for senescence based on its gene expression profile. The details of the DeepScence model are described below.

#### Input data

DeepScence takes as input a properly filtered expression count matrix. Data filtering and quality control can be performed using standard processing pipelines such as Seurat or Scanpy[58, 59]. This step is essential for removing low-quality cells that may affect the performance of DeepScence, as DeepScence does not perform any additional filtering. The performance of DeepScence is robust across varying data filtering stringencies. For example, we reprocessed the data using an alternative filtering criterion, in which cells with fewer than 200, more than 7000 expressed genes, and more than 20% mitochondrial reads were removed from all scRNA-seq datasets. DeepScence maintained comparable performance and remained among the top-performing methods (Supplementary Figures 3–4).

The filtered matrix is first denoised using DCA (version 0.3.1) [28] with default parameters. The denoised expression count matrix is then library size normalized and log-transformed using the `scanpy` (version 1.9.8) functions `sc.pp.normalize_total` and `sc.pp.log1p` with default parameters. The log-normalized expression matrix is then subsetted to retain only the genes in the CoreScence gene set. Finally, the subsetted expression matrix is scaled using the `scanpy sc.pp.scale` function, so that the gene expression values have zero mean and unit standard deviation for each gene across all cells.

#### Architecture

Let ***X*** denote the denoised and subsetted gene expression count matrix, and let ***X*** ^***′***^ denote the denoised, log-normalized, subsetted, and scaled expression matrix. DeepScence employs an autoencoder architecture similar to DCA [28]. Specifically, ***X*** ^***′***^ is encoded into a bottleneck layer consisting of two nodes, and then decoded into three output layers. Each output layer matches the size of the input ***X*** ^***′***^ and represents the estimated dropout, mean, and dispersion parameters of the ZINB distribution. Since the input ***X*** ^***′***^ is normalized by library size, the encoded 2-dimensional representation, as well as the estimated dropout, mean, and dispersion matrices, are not affected by library sizes. The model can be specified in the formulation below.

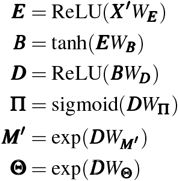

Here, ***E*** represents the encoder layer with 32 neurons, ***B*** is the bottleneck layer with 2 neurons, and ***D*** is the decoder layer with 32 neurons. **Π, *M***^***′***^, and **Θ** are the output dropout, mean, and dispersion corresponding to the ZINB distribution. 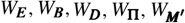, and *W*_**Θ**_ are the parameters to be estimated during the model fitting process.

#### Objective function

The objective function of the model above, which will be minimized during the model fitting process, is defined as:

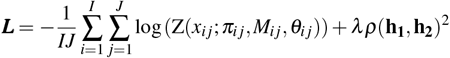

Here, Z represents the probability density function of the ZINB distribution. *I* and *J* represent the total number of CoreScence genes and cells, respectively. *x*_*ij*_, *π*_*ij*_, *M*_*ij*_, and *θ*_*ij*_ represent the gene expression count, dropout, mean, and dispersion for *i*th gene and *j*th cell, respectively. **h**_1_ and **h**_2_ are two vectors representing the values of the two neurons in the bottleneck layer, before tanh activation. *ρ*(**h**_**1**_, **h**_**2**_) represents the Pearson correlation coefficient between **h**_1_ and **h**_2_.

The objective function of DeepScence consists of two components. The first component is the negative log-likelihood of the ZINB distribution. Minimizing this term ensures that the autoencoder learns essential information from the input gene expression count matrix. Before calculating this term, note that the estimated mean matrix, ***M***^***′***^, is converted to ***M*** by multiplying it by the size factors. The second component is the Pearson correlation coefficient between two neurons in ***H***. Minimizing this term ensures that the two neurons in ***H*** are almost uncorrelated, allowing them to capture information related to senescence and unrelated information, respectively.

The objective function is a weighted average of the two components, controlled by a positive hyperparameter *λ*. A *λ* value that is too low may cause the two neurons in the bottleneck layer to become correlated, diluting the senescence-related information captured by each neuron. Conversely, a *λ* value that is too high may lead the model to overly focus on minimizing correlation, neglecting the ZINB component that learns from the data. We empirically observed the model’s behavior under different *λ* values and found that *λ* = 1 consistently results in a low absolute Pearson correlation (|*ρ*| < 0.1), while still generating desirable outcomes. Therefore, we set the default value of *λ* to 1, although users can adjust this value if needed.

#### Model training

By default, DeepScence is trained for 300 epochs without mini-batching at a learning rate of 0.005 using the Adam optimizer[60]. During each epoch, 10% of the cells are used as a validation set, and the validation loss is recorded based on the ZINB loss for these cells. An early stopping mechanism with learning rate reduction is implemented: if the validation loss does not improve for 10 consecutive epochs, the learning rate is halved. If there is no improvement for 30 epochs, the entire training process is stopped.

#### Output

After model fitting, DeepScence outputs the values of the neuron responsible for capturing senescence information in the bottleneck layer as continuous senescence scores. Since the roles of the two neurons are undetermined during model fitting, DeepScence uses a *post hoc* approach to identify the neuron that captures senescence information. Specifically, for each of the two neurons, DeepScence calculates the absolute value of the Pearson correlation coefficient between the neuron’s values and the expression of each gene in CoreScence, based on ***X*** ^***′***^. The neuron with the higher average absolute correlation across all CoreScence genes is identified as the one capturing senescence information.

Finally, to ensure that the output senescence score is positively correlated with the cells’ senescence levels, DeepScence calculates the Pearson correlation between the output score and the expression of the CDKN1A gene, a canonical marker of senescence. If the correlation is negative, the sign of the output score is flipped.

#### Binarization of the continuous score

Cellular senescence, a continuous process by nature [61], is best characterized by a continuous score, such as that provided by DeepScence. However, DeepScence also offers the option to convert the continuous senescence score into a binary score, which can be useful for certain downstream analyses, such as comparing the proportion of SnCs across samples or cell types. The binary score is obtained through a two-step process.

In the first step, an uncertainty score is calculated for each cell by comparing its continuous senescence score to a null distribution. Specifically, for each cell in the expression matrix ***X*** ^***′***^, the gene expression values are randomly permuted across all CoreScence genes. The permuted gene expression values are then input into the DeepScence autoencoder, already fitted with the original data, and the value of the neuron that captures senescence information is recalculated. This process is repeated 50 times to generate 50 permuted scores. The cell’s uncertainty score is defined as the proportion of times the permuted score exceeds the cell’s original DeepScence score. A senescence score cutoff is determined as the smallest value such that less than 1% of cells with senescence scores above the cutoff have uncertainty scores greater than 0.5.

In the second step, a mixture of normal distributions is fitted to the continuous senescence scores of all cells using the `GaussianMixture` function from the Python `sklearn` package (version 1.3.2). The number of normal components is selected from 2 to 10 based on the lowest Akaike Information Criterion (AIC). Cells are assigned to the normal components according to posterior probabilities. Among the normal distributions whose 5% quantiles are greater than the senescence score cutoff defined in the previous step, the distribution with the smallest mean is selected. Senescent cells are defined as those assigned to this distribution or any distribution with a mean larger than that of the selected distribution.

### Competing methods

#### Single marker gene approach

For the scRNA-seq dataset, the gene expression count matrix was normalized by library size and then log-transformed using the functions `sc.pp.normalize_total` and `sc.pp.log1p` with default settings from the Python package `scanpy`. The log-normalized expression values of individual genes, such as CDKN1A and CDKN2A, were used as continuous senescence scores, with cells exhibiting higher expression levels being more likely to be senescent. For binarization, cells with positive gene expression values were classified as senescent, while cells with zero expression values were classified as non-senescent.

#### Gene ranking approach based on SnGs

For the scRNA-seq dataset, the log-normalized gene expression matrix was obtained as described above. Using CoreScence or each of the published SnGs as input, we then calculated continuous senescence scores with the `gsva` function and the ssGSEA method in the R package `GSVA` (version 1.52.0), as well as the `AUCell_run` function in the R package `AUCell` (version 1.26.0). Note that for SnGs with directional information, only genes that induce senescence were used as input.

`GSVA` and `AUCell` do not provide functions to binarize the continuous senescence scores. However, the original SenMayo study [20], which uses the ssGSEA method, treats the top 10% of cells with the highest senescence scores as senescent cells. Therefore, we only evaluated the binarized scores for the SenMayo gene set using the ssGSEA method. The top 10% of cells with the highest senescence scores were predicted as senescent cells, while the remaining cells were predicted as non-senescent.

#### SenCID

For each scRNA-seq dataset, the Python package SenCID (version 1.0.0) was used to obtain both continuous and binarized senescence scores with default settings, using the gene expression count matrix as input.

#### DeepScence with alternative gene sets

We replaced CoreScence with one of the six published gene sets containing more than 100 genes (SenSig, SenMayo, CellAge, GenAge, CSGene, and SeneQuest) and re-ran DeepScence on each scRNA-seq and ST dataset. All other settings, including the autoencoder architecture and training details, remained the same.

### Evaluation of SnC identification methods using datasets with established gold standards

#### Collection of in vitro scRNA-seq datasets

We collected *in vitro* scRNA-seq datasets from six studies that profiled both senescent and non-senescent cells.

In the first study [27], the gene expression count matrix for the WI-38 cell line was downloaded from GEO with accession GSE175533. The downloaded matrix had already been filtered, and no additional cell filtering was performed. Cells with the highest population doubling (PDL = 50) were considered senescent cells (SnCs), while negative controls (hTERT) were considered non-senescent.

In the second study [32], the gene expression count matrix for the HCA2 cell line was downloaded from GEO with accession GSE119807. Following the same cell filtering criteria as in [32], cells with more than 500 detected genes and a mitochondrial percentage of less than 10% were included. Cells labeled as “senescence” in the metadata provided by the original study were considered SnCs, while those labeled as ‘LowPDCtrl’ were considered non-senescent.

In the third study [34], the gene expression count matrix for the IMR-90 cell line was downloaded from GEO with accession GSE115301. Following the same cell filtering criteria as in [34], cells with less than 15% of reads mapped to mitochondrial genes and with positive expression in at least 2,500 genes were retained. Cells labeled as “RIS” (radiation-induced senescence) in the downloaded metadata were considered SnCs, while those labeled as “Growing” were considered non-senescent.

In the fourth study [35], the gene expression count matrix for the second IMR-90 cell line was downloaded from GEO with accession GSE94980. The downloaded matrix had already been filtered, and no additional cell filtering was performed. Cells infected with the reprogramming factors OSKM (indicated by “OSKM” in cell names) were considered SnCs, while the controls (indicated by “Vector” in cell names) were treated as non-senescent.

In the fifth study [33], the gene expression count matrix for the HUVEC cell line was downloaded from GEO with accession GSE102090. The downloaded matrix had already been filtered, and no additional cell filtering was performed. As detailed in the original publication, cells with names ending in the suffix “-2”, indicating the second batch, were considered senescent SnCs, while cells from the first batch were considered non-senescent.

In the sixth study[36], the gene expression count matrix for the WI-38 cell line was downloaded from GEO under accession GSE226225. Following the same cell filtering criteria as in the original study, cells with less than 12% of reads mapped to mitochondrial genes and with between 1,000 and 120,000 total positive expression counts were retained. Cells labeled as “CTRL” were considered non-senescent, while the remaining cells were classified as SnCs.

For datasets with more senescent cells than non-senescent cells, which is unlikely in *in vivo* settings, we randomly removed SnCs to equalize the numbers of SnCs and non-senescent cells. Finally, we retained genes with non-zero expression in at least 1% of the cells in each dataset.

#### Collection of in vivo scRNA-seq datasets

We collected eight *in vivo* scRNA-seq datasets that profiled both cells from diseased and normal conditions. We included only cell types for which immunohistochemistry validation, such as *β*-galactosidase and p16 staining, was performed to confirm that SnCs were more enriched in cells from diseased conditions.

In the first study [37], the gene expression count matrix and cell type annotations for mouse testis cells were downloaded from GEO with accession GSE183625. We included only Leydig cells, in which *β*-gal staining was performed in the original study. Following the same cell filtering criteria as in [37], cells with less than 20% of reads mapped to mitochondrial genes and with positive expression in 200 to 7,000 genes were retained. Gene names were converted to their human homologs using the conversion table available on the Mouse Genome Informatics (MGI) website [62]. Cells labeled as “O50t” in the metadata were considered to be in the SnC-enriched condition, while cells labeled as “Nt” represented the normal condition.

In the second study [38], the gene expression count matrix and cell type annotations for mouse muscle cells were downloaded from GEO with accession GSE214892. The matrix had already been filtered, and no additional cell filtering was performed. All cell types were included in this study. Gene names were converted to their human homologs using the conversion table available on the Mouse Genome Informatics (MGI) website [62]. Cells labeled as “CTX” in the metadata were considered to be in the SnC-enriched conditions, while cells labeled as “CTRL” represented the normal condition.

In the third study [39], the gene expression count matrix and cell type annotations for human lung cells were downloaded from GEO with accession GSE190889. We included only AT1 cells, in which *β*-gal staining was performed in the original study. The matrix had already been filtered, and no additional cell filtering was performed. Cells labeled as “IPF” in the metadata were considered to be in the SnC-enriched conditions, while cells labeled as “healthy” represented the normal condition.

In the fourth study [63], the gene expression count matrix for human oral mucosa cells was downloaded from GEO with accession GSE164241. Following the same cell filtering criteria as in [63], cells with less than 10% of reads mapped to mitochondrial genes, positive expression in 200 to 5,000 genes, and a total number of reads between 1,000 and 25,000 were retained. The original study did not provide cell type annotations. To perform cell type annotation, we first applied log normalization to the filtered expression count matrix using the Seurat function `NormalizeData`. We then ran PCA using the Seurat functions `FindVariableFeatures`, `ScaleData`, and `RunPCA` with default parameters. Cell clustering was performed using the Seurat function `FindNeighbors` with the top 15 PCs, followed by `FindClusters` with a resolution of 0.8. Cell type annotation was then performed using the R package GPTCelltype [64], with the tissue name set to ‘human oral mucosa’ and the model set to ‘gpt-4’. We included only fibroblasts, in which *β*-gal staining was performed in the original study. Cells labeled as “periodontitis” in the metadata were considered to be in the SnC-enriched conditions, while cells labeled as “healthy” represented the normal condition.

In the fifth study [42], the gene expression count matrix for cardiomyocytes and non-cardiomyocyte cells from mouse heart tissue was downloaded from GEO with accession GSE221744. Following the same cell filtering criteria as in [42], cardiomyocytes with more than 4,000 detected genes per cell were retained. Non-cardiomyocytes with more than 400 and fewer than 5,000 detected genes, more than 5,000 total counts, and less than 30% mitochondrial gene content were retained. For non-cardiomyocytes, unsupervised clustering was performed using the Seurat function FindNeighbors with the top 20 principal components, followed by FindClusters with a resolution of 0.3. Cell type annotation was carried out using the R package GPTCelltype[64], with the tissue name set to mouse heart and the model set to gpt-4. We included only endothelial cells (ECs), in which p16 staining was performed in the original study. Cells labeled as “TAC2w” in the metadata were considered to be in the SnC-enriched conditions, while cells labeled as “Sham” represented the normal condition. Notice that data from cardiomyocytes and non-cardiomyocytes were generated using different sequencing platforms, so they were treated as two separate datasets.

In the sixth study [41], the filtered gene expression count matrix was downloaded directly from GEO with accession GSE229553, so no additional filtering was performed. Unsupervised clustering was carried out using Seurat’s FindNeighbors function with the top 20 principal components, followed by FindClusters with a resolution of 0.8. Cell type annotation was performed using the R package GPTCelltype [64], with the tissue name set to mouse brain and the model set to gpt-4. We included only microglia, based on literature evidence that microglia are prone to senescence in Alzheimer’s disease (AD) tauopathy [45], in which p16, p21, GLB1, and *γ*H2AX staining were performed. Cells labeled as “tau” in the metadata were considered to be in the SnC-enriched conditions, while cells labeled as “wt” represented the normal condition.

In the seventh study [43], the filtered and preprocessed expression matrix was directly downloaded from GEO with accession GSE203447, so no additional filtering was performed. Cell type annotations were provided in the downloaded data. All cell types were used in this dataset, as senescent cells (SnCs) were identified by FACS using the p16-FDR mouse line. Cells with mCherry.Count > 0 in the metadata were considered SnCs, while cells with mCherry.Count = 0 were considered non-SnCs.

Finally, for all datasets, we retained genes with positive expression in at least 1% of the retained cells.

#### Standard filtering procedure

To demonstrate DeepScence’s robustness under different filtering stringencies, we downloaded the unfiltered gene expression count matrices of all scRNA-seq datasets from the original studies. We then performed an in-house standardized filtering, retaining cells with more than 200 and fewer than 7,000 expressed genes, and less than 20% mitochondrial gene content. DeepScence and other competing methods were subsequently applied to the datasets that underwent this standardized filtering.

#### Collection of spatial transcriptomics datasets

We downloaded 10x Visium ST datasets of mouse muscle tissues collected 2 and 5 days post-notexin-injury [44]. The gene expression count matrix, along with the annotation indicating whether a spatial spot was in the injured or healthy regions, was downloaded from Dryad (DOI: 10.5061/dryad.t4b8gtj34). The matrix had already been filtered, so no additional cell filtering was performed. Gene names were converted to their human homologs using the conversion table available on the Mouse Genome Informatics (MGI) website [62]. We also generated zoomed-in images of cellular morphology using the accompanying H&E images.

We downloaded STOmics Stereo-seq ST datasets of human brains from GEO with accession GSE269906 [65]. These datasets provide paired samples from individuals with Alzheimer’s disease (AD) and their normal aging counterparts. For our analysis, we included the two samples classified as “severe AD” along with their matched normally aging controls. Since the Stereo-seq platform [66] offers subcellular resolution, we applied a cell-level analysis pipeline analogous to single-cell workflows, following the instructions of the Python package stereopy. Specifically, we set *n*_*neighbors*_=10 and *n*_*pcs*_=40 to compute the cell neighborhood graph, and used the Leiden algorithm for unsupervised clustering with a resolution of 0.7. Prior studies [41, 45] have reported that microglial senescence is more prevalent in AD compared to non-diseased brains, supported by increased immunostaining of senescence markers such as *β*-galactosidase, p16, p21, and *γ*H2AX. Therefore, we used the canonical microglial marker gene CSF1R [67] to identify microglial clusters and included only microglial cells for benchmarking SnC identification methods.

#### Generation of simulated 10x Xenium data

We downloaded the ‘Human Multi-Tissue and Cancer Panel’ from the 10x pre-designed panels, which includes 377 human genes. Since the original panel contained very few senescence-associated genes, we expanded it by adding randomly selected senescence-associated genes. For DeepScence, the genes were randomly selected from CoreScence, and for SenCID, the genes were randomly selected from the list of genes used in SenCID’s SVM model. For each cell line in the *in vitro* studies, we then subsetted the expression count matrix to include only the genes from the expanded panel.

#### Evaluation details

To evaluate the continuous senescence scores, ROC curves and AUROC were generated using the R package `pROC` (version 1.18.5).

To evaluate the binarized senescence scores, we used the functions `accuracy_score` and `f1_score` from the Python package `scikit-learn` (version 1.1.3). Let TP, TN, FP, and FN represent the true positives, true negatives, false positives, and false negatives, respectively. The accuracy is defined as:

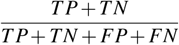

The F1 score is defined as:

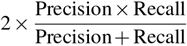

where

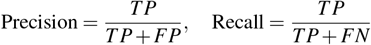

### Analysis of normally aging datasets

#### Collection of scRNA-seq datasets

We collected scRNA-seq data from the Tabula Sapiens study [49]. The filtered and preprocessed expression matrix was downloaded from Figshare at https://figshare.com/articles/dataset/Tabula_Sapiens_v2/27921984. We obtained cell metadata including sample ages, tissue names, and cell type ontology classes annotated by the original study. To achieve a comparable level of granularity in cell type annotations across all tissues, we manually re-annotated the cell types, as detailed in Supplementary Table S3.

We classified samples into age groups using the median age across all samples, which was 59 years, as the threshold. Samples with age ≥ 59 years were assigned to the old group, and those with age *<* 59 years to the young group.

We filtered out cell types that were present in fewer than three tissues, as well as tissues that contained fewer than three cell types. Tissue–cell type pairs with an average log-normalized expression of CDKN1A less than 0.05 were also removed, to ensure the presence of a certain level of SnCs in the data. Tissue–cell type pairs with fewer than 300 cells in either the old or young age group were also filtered out. After filtering, a total of nine cell types across 17 tissues remained.

#### Collection of spatial transcriptomics datasets

We downloaded the 10x Visium ST dataset of normally aging mouse mammary tissues [50] from GEO with accession GSE216542. The dataset includes two 3-month-old samples and two 18-month-old samples. The expression matrix had already been filtered, so no additional cell filtering was performed. Spatial spots from 18-month-old mice were assigned to the old group, and spots from 3-month-old mice to the young group.

#### Evaluation details

For each tissue–cell type pair in the TS dataset, we defined cells with the top 1% of DeepScence scores as top quantile cells, and the remaining cells as bottom quantile cells. Let *a* denote the number of top quantile cells from the old group, *b* the number from the young group, *c* the number of bottom quantile cells from the old group, and *d* the number from the young group. We then calculated the SnC enrichment as:

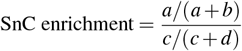

SnC enrichment for other competing methods and for the mouse mammary tissue ST dataset was computed in a similar manner.

### Computational Efficiency

A total of 100, 500, 1,000, 5,000, 10,000, 50,000, and 100,000 cells were randomly sampled from the Tabula Sapiens dataset after pooling cells across all tissues and cell types. DeepScence was executed on each subset five times. The running time and peak memory usage were recorded using the Python `time` and `memory_profiler` packages. The runtime and peak memory usage were averaged across the five repeats for each cell count.

## Acknowledgments

Y.Q., C.C., J.X., C.G., X.W., A.N., and Z.J. were funded by the National Institutes of Health under award number U54AG075936. Y.Q. and Z.J. were funded by the National Institutes of Health under award number R35GM154865. R.D. and L.G. were funded by the National Institutes of Health under award number UH3CA268096.

## Author contributions

All authors contributed to the conceptualization of the study. Y.Q. developed the DeepScence software. Y.Q. and B.J. performed the analysis. Y.Q. and Z.J. jointly wrote the manuscript.

## Competing interests

A.N. receives funding from Genentech, Genmab, MedImmune/AstraZeneca, and Seattle Genetics. A.N. is a consultant of Sanofi, Promega, and Leap Therapeutics.

## Code availability

The DeepScence Python package is freely available on GitHub: https://github.com/anthony-qu/DeepScence. The analysis code for reproducing the results can also be accessed on GitHub: https://github.com/anthony-qu/DeepScence_analysis.

## Data availability

All data analyzed in this study were obtained from previously published studies.

Information on differentially expressed genes from the two bulk RNA-seq studies was downloaded from: https://www.ncbi.nlm.nih.gov/pmc/articles/PMC6698740/bin/gkz555_supplemental_files.zip and https://cdn.elifesciences.org/articles/70283/elife-70283-fig1-data2-v3.xlsx.

The scRNA-seq data with *in vitro* senescence inductions were downloaded from GEO. Specifically, the WI-38 dataset was downloaded from https://www.ncbi.nlm.nih.gov/geo/query/acc.cgi?acc=GSE175533, the HCA2 dataset from https://www.ncbi.nlm.nih.gov/geo/query/acc.cgi?acc=GSE119807, the two IMR-90 datasets from https://www.ncbi.nlm.nih.gov/geo/query/acc.cgi?acc=GSE115301 and https://www.ncbi.nlm.nih.gov/geo/query/acc.cgi?acc=GSE94980, and the HUVEC dataset from https://www.ncbi.nlm.nih.gov/geo/query/acc.cgi?acc=GSE102090.

The *in vivo* scRNA-seq data were downloaded from GEO. Specifically, the mouse testes EAO dataset was downloaded from https://www.ncbi.nlm.nih.gov/geo/query/acc.cgi?acc=GSE183625, the mouse muscle injury data from https://www.ncbi.nlm.nih.gov/geo/query/acc.cgi?acc=GSE214892, the human lung IPF data from https://www.ncbi.nlm.nih.gov/geo/query/acc.cgi?acc=GSE190889, and the human oral mucosa data from https://www.ncbi.nlm.nih.gov/geo/query/acc.cgi?acc=GSE164241. The mouse heart dataset was downloaded from https://www.ncbi.nlm.nih.gov/geo/query/acc.cgi?acc=GSE221744, the mouse brain AD dataset was downloaded from https://www.ncbi.nlm.nih.gov/geo/query/acc.cgi?acc=GSE229553, the mouse lung cancer dataset was downloaded from https://www.ncbi.nlm.nih.gov/geo/query/acc.cgi?acc=GSE203447

The 10x Visium ST dataset of mouse muscle injury by notexin injection was downloaded from https://datadryad.org/stash/dataset/doi:10.5061/dryad.t4b8gtj34. The Human brain stereo-seq dataset was downloaded from https://www.ncbi.nlm.nih.gov/geo/query/acc.cgi?acc=GSE269906

## Notes

### Competing Interest Statement

A.N. receives funding from Genentech, Genmab, MedImmune/AstraZeneca, and Seattle Genetics. A.N. is a consult of Sanofi, Promega, and Leap Therapeutics.

### Summary of Updates

We have added new analysis and benchmark results in the revised manuscript.

