## Supplementary Figure for "Single-cell and spatial detection of senescent cells using DeepScence"

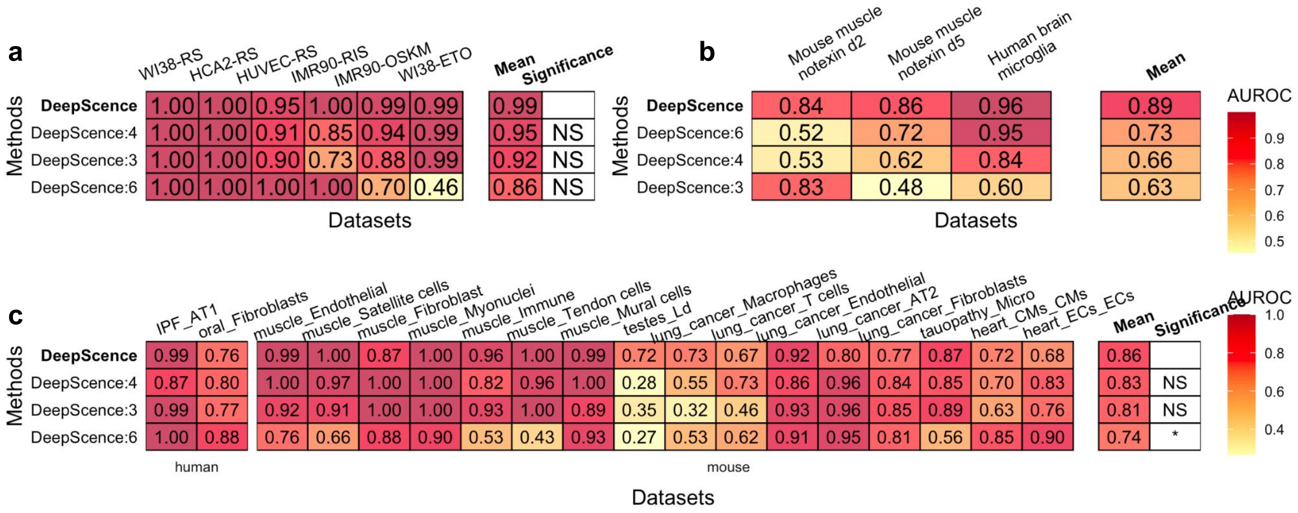


| Methods | WI38-RS | HCA2-RS | HUVEC-RS | IMR90-RIS | IMR90-OSKM | WI38-ETO | Mean | Significance |
|---|---|---|---|---|---|---|---|---|
| **DeepScence** | 1.00 | 1.00 | 0.95 | 1.00 | 0.99 | 0.99 | 0.99 | |
| DeepScence:4 | 1.00 | 1.00 | 0.91 | 0.85 | 0.94 | 0.99 | 0.95 | NS |
| DeepScence:3 | 1.00 | 1.00 | 0.90 | 0.73 | 0.88 | 0.99 | 0.92 | NS |
| DeepScence:6 | 1.00 | 1.00 | 1.00 | 1.00 | 0.70 | 0.46 | 0.86 | NS |

| Methods | Mouse muscle notexin d2 | Mouse muscle notexin d5 | Human brain microglia | Mean |
|---|---|---|---|---|
| **DeepScence** | 0.84 | 0.86 | 0.96 | 0.89 |
| DeepScence:6 | 0.52 | 0.72 | 0.95 | 0.73 |
| DeepScence:4 | 0.53 | 0.62 | 0.84 | 0.66 |
| DeepScence:3 | 0.83 | 0.48 | 0.60 | 0.63 |

| Methods | IPF_AT1 | oral_Fibroblasts | muscle_Endothelial | muscle_Satellite cells | muscle_Fibroblast | muscle_Myonuclei | muscle_Immune | muscle_Tendon cells | muscle_Mural cells | testes_Ld | lung_cancer_Macrophages | lung_cancer_T cells | lung_cancer_Endothelial | lung_cancer_AT2 | lung_cancer_Fibroblasts | tauopathy_Micro | heart_CMs_CMs | heart_ECs_ECs | Mean | Significance |
|---|---|---|---|---|---|---|---|---|---|---|---|---|---|---|---|---|---|---|---|---|
| **DeepScence** | 0.99 | 0.76 | 0.99 | 1.00 | 0.87 | 1.00 | 0.96 | 1.00 | 0.99 | 0.72 | 0.73 | 0.67 | 0.92 | 0.80 | 0.77 | 0.87 | 0.72 | 0.68 | 0.86 | |
| DeepScence:4 | 0.87 | 0.80 | 1.00 | 0.97 | 1.00 | 1.00 | 0.82 | 0.96 | 1.00 | 0.28 | 0.55 | 0.73 | 0.86 | 0.96 | 0.84 | 0.85 | 0.70 | 0.83 | 0.83 | NS |
| DeepScence:3 | 0.99 | 0.77 | 0.92 | 0.91 | 1.00 | 1.00 | 0.93 | 1.00 | 0.89 | 0.35 | 0.32 | 0.46 | 0.93 | 0.96 | 0.85 | 0.89 | 0.63 | 0.76 | 0.81 | NS |
| DeepScence:6 | 1.00 | 0.88 | 0.76 | 0.66 | 0.88 | 0.90 | 0.53 | 0.43 | 0.93 | 0.27 | 0.53 | 0.62 | 0.91 | 0.95 | 0.81 | 0.56 | 0.85 | 0.90 | 0.74 | * |

**Supplementary Figure 1.** AUROCs for DeepScence using different thresholds for constructing CoreScence across *in vitro* scRNA-seq datasets (**a**), ST datasets (**b**), and *in vivo* scRNA-seq datasets (**c**). Methods are ordered in decreasing order of average AUROC. Paired t-tests were conducted to compare the performance between default DeepScence and its variants. "*" indicates 0.01 < p-value < 0.05, and "NS" indicates not significant (p-value > 0.05).

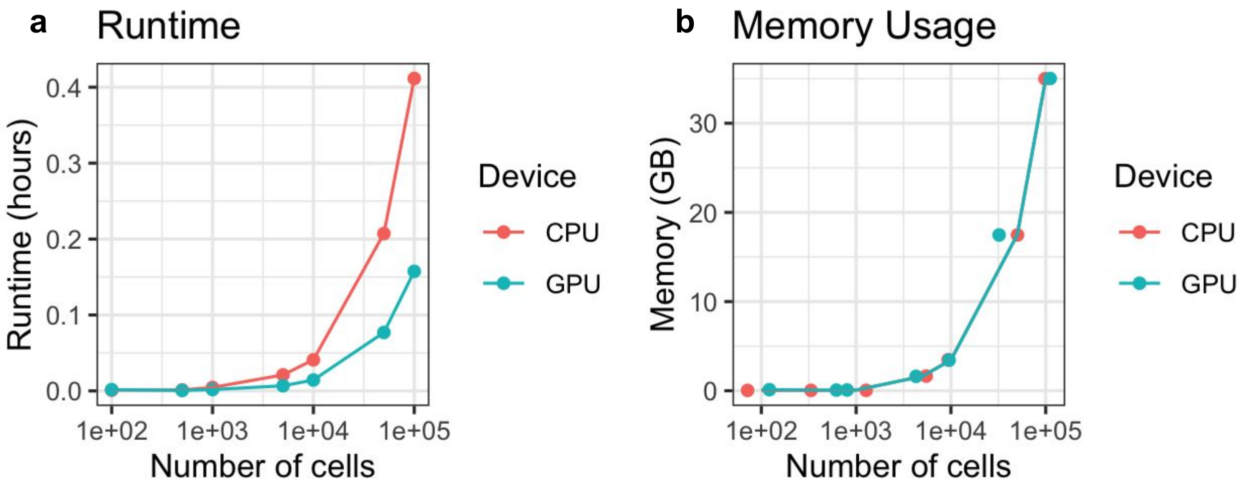


**Supplementary Figure 2.** Runtime (**a**) and peak memory usage (**b**) for executing DeepScence with different numbers of cells in the input data.

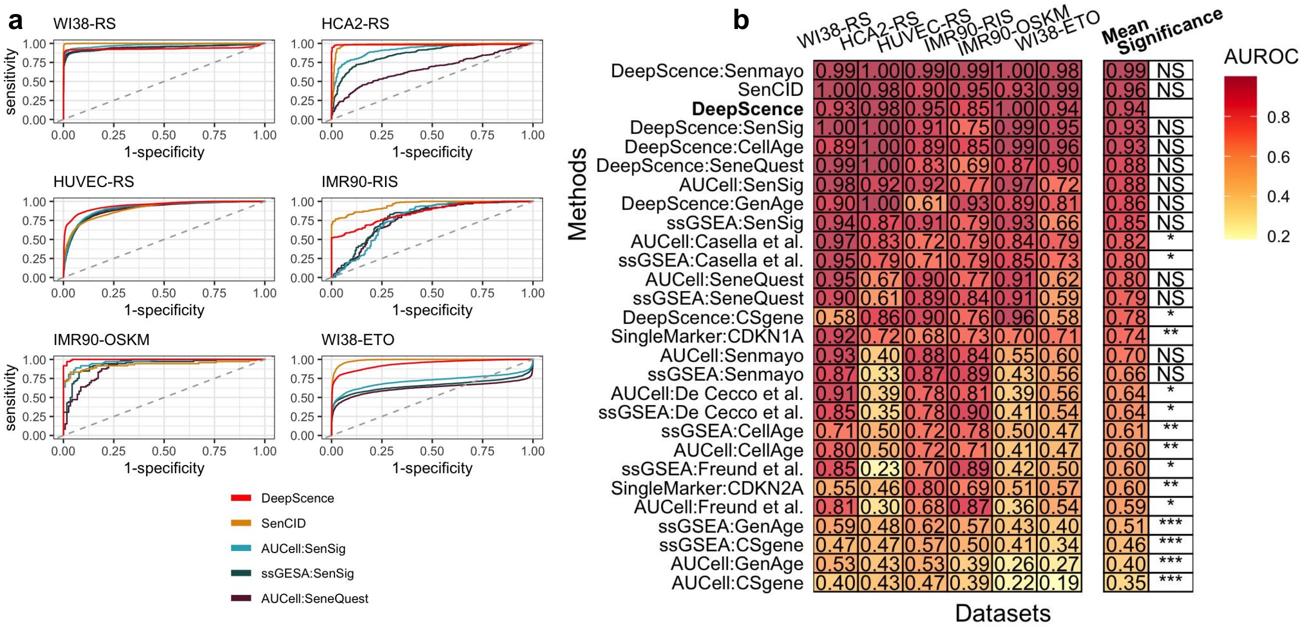


**Supplementary Figure 3. a,** AUC curves for the top 5 performing methods among *in vitro* datasets under standard data preprocessing pipeline. **b,** AUROCs for all methods among *in vitro* datasets under standard data preprocessing pipeline. Methods are ordered in decreasing order by average AUROCs. Paired t-test was conducted to compare the performance between DeepScence and each of the other methods. “***” indicates p-value < 0.001, "**" indicates 0.001 < p-value < 0.01, "*" indicates 0.01 < p-value < 0.05, and "NS" indicates not significant (p-value > 0.05).

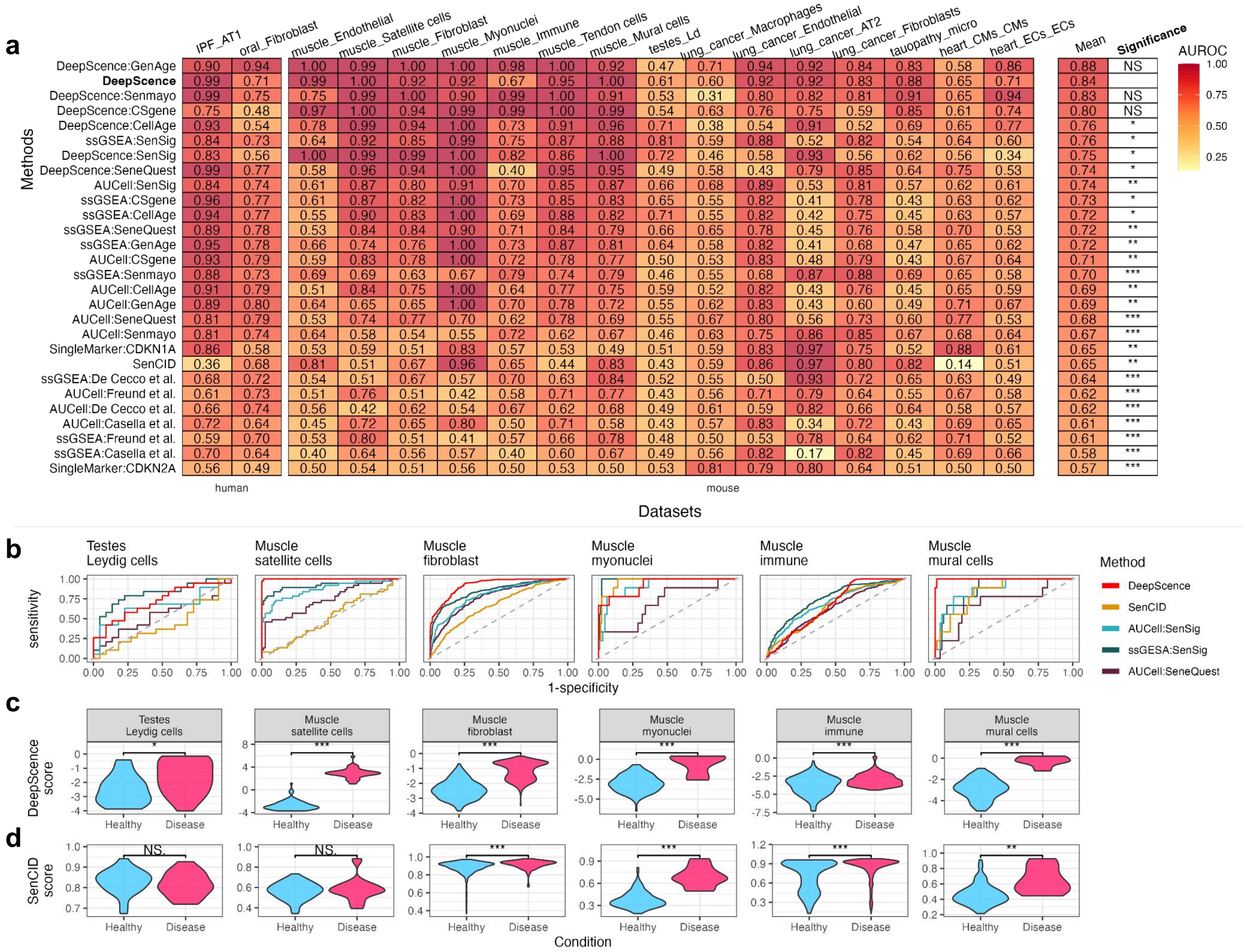


**Supplementary Figure 4. a,** AUROCs for all methods across *in vivo* datasets under standard preprocessing pipeline. Methods are ordered in decreasing order by average AUROCs. Paired t-test was conducted to compare the performance between DeepScence and each of the other methods. “***” indicates p-value < 0.001, "**" indicates 0.001 < p-value < 0.01, "*" indicates 0.01 < p-value < 0.05, and "NS" indicates not significant (p-value > 0.05). **b,** ROC curves for six example cell types in *in vivo* datasets under standard preprocessing, showing methods with top performance in *in vitro* datasets. **c,** Distribution of SenCID and DeepScence scores under standard preprocessing, comparing cells from healthy and diseased conditions. Wilcoxon test was conducted to compare the two distributions in each case. ’*’ indicates p-value between 0.01 and 0.05, ’**’ indicates p-value between 0.001 and 0.01, and ’***’ indicates p-value < 0.001.
